# Crystal structure of a *Thermus aquaticus* diversity-generating retroelement variable protein

**DOI:** 10.1101/432187

**Authors:** Sumit Handa, Kharissa L Shaw, Partho Ghosh

## Abstract

Diversity-generating retroelements (DGRs) are widely distributed in bacteria, archaea, and microbial viruses, and bring about unparalleled levels of sequence variation in target proteins. While DGR variable proteins share low sequence identity, the structures of several such proteins have revealed the C-type lectin (CLec)-fold as a conserved scaffold for accommodating massive sequence variation. This conservation has led to the suggestion that the CLec-fold may be useful in molecular surface display applications. Thermostability is an attractive feature in such applications, and thus we studied the variable protein of a DGR encoded by the thermophile *Thermus aquaticus*. We report here the 2.8 Å resolution crystal structure of the variable protein from the *T. aquaticus* DGR, called TaqVP, and confirm that it has a CLec-fold. Remarkably, its variable region is nearly identical in structure to those of several other CLec-fold DGR variable proteins despite low sequence identity among these. TaqVP was found to be thermostable, which appears to be a property shared by several CLec-fold DGR variable proteins. These results provide impetus for the pursuit of the DGR variable protein CLec-fold in molecular display applications.

## Introduction

Diversity generating-retroelements (DGRs) are unique and unparalleled generators of massive protein sequence diversity [1, 2]. At least 10^20^ sequences are possible in proteins diversified by these retroelements [3]. This scale of variation exceeds by several orders of magnitude the variation brought about by the adaptive immune systems of jawed and jawless vertebrates, the only other examples of natural massive protein sequence variation [4, 5]. In these immune systems, massive sequence variation of immunoreceptors permits the recognition of novel ligands and is a robust means for adaptation to dynamic environments. Similarly, massive sequence variation by DGRs appears to enable adaptation to dynamic environments for the ecologically diverse microbes – bacteria, archaea, and microbial viruses – that encode them. These include constituents of the human microbiome [2, 3, 6–12] as well as of the microbial ‘dark matter’ which constitutes a major fraction of microbial life [13–17].

Three fundamental components define DGRs: a reverse transcriptase (RT) that has a unique sequence motif [1, 2, 18], a variable region (*VR*) that forms part of the coding sequence of a DGR diversified protein, and a template region (*TR) that is nearly identical to VR and is located proximally to VR* (Fig 1). The *TR* serves as an invariant store of amino acid coding information. This information is transferred from *TR* to VR, and during this process adenines within *TR* are specifically mutated to other bases. A recent study on the prototypical DGR of *Bordetella* bacteriophage indicates that the DGR RT in association with a second DGR protein, the accessory variability determinant (Avd), is necessary and sufficient for mutagenesis of adenines in *TR* [19]. Adenine-specific mutagenesis culminates in substitutions occurring only at amino acids that have adenines in their codons in *TR*. AAY is the most recurrent adenine-containing codon in the *TR* of a variety of DGRs [2]. As previously noted, adenine-mutagenesis of Asn-encoding AAY can result in the encoding of 14 other amino acids, which cover the gamut of amino acid chemical character, but cannot result in a stop codon [20].

DGR variable proteins are divergent in sequence (roughly ≤ 20% identity) [2, 20]. To date, structures of three DGR variable proteins have been determined — *Bordetella* bacteriophage Mtd [20, 21], Treponema denticola TvpA [3], and *Nanoarchaeota* AvpA [22]. Despite the sequence divergence of these proteins, they all share a C-type lectin (CLec)-fold, which appears to be an evolutionarily conserved scaffold among DGRs for accommodating massive protein sequence variation. The CLec-fold, despite its nomenclature, is a general ligand-binding fold [23]. Variable amino acids in these CLec-fold DGR variable proteins are solvent-exposed and form ligand-binding sites. Some DGR variable proteins, including those that exist in the human virome [6], are predicted instead to have an immunoglobulin (Ig)-fold as a scaffold for variation. The β-strand body of the Ig-fold appears to be the locus for variation in these proteins, rather than loop regions as in antibodies and T cell receptors. Yet other DGR variable proteins have folds that at present cannot be assigned by sequence alone to CLec- or Ig-folds, or to other protein folds [2].

DGR variable proteins with characterized CLec-folds have the potential to add to the existing toolbox of selectable variable protein scaffolds for molecular display purposes, which includes affibodies, fibronectin type III (FN3) domains, and designed ankyrin repeat proteins (DARPins) among others [24, 25]. As thermostability is an attractive property of selectable variable proteins, we focused on a DGR of the thermophile *Thermus aquaticus* [2]. The *T. aquaticus* DGR has a genetic structure similar to that of that of the *Bordetella* bacteriophage DGR (Fig. 1). The variable protein of the *T. aquaticus* DGR, which we call TaqVP, is predicted to have a CLec-fold [2]. While a common theme in DGR variable proteins is surface-display [12], TaqVP appears to lack a surface-targeting signal sequence, suggesting that sequence variation in the protein may have an intracellular function instead. We determined the 2.8 Å resolution crystal structure of TaqVP and confirmed that it indeed has a CLec-fold. We also characterized the thermostability of TaqVP, along with those of the previously structurally characterized DGR variable proteins Mtd and TvpA, and found that thermostability was a shared property of several DGR CLec-fold proteins.

**Figure 1.**
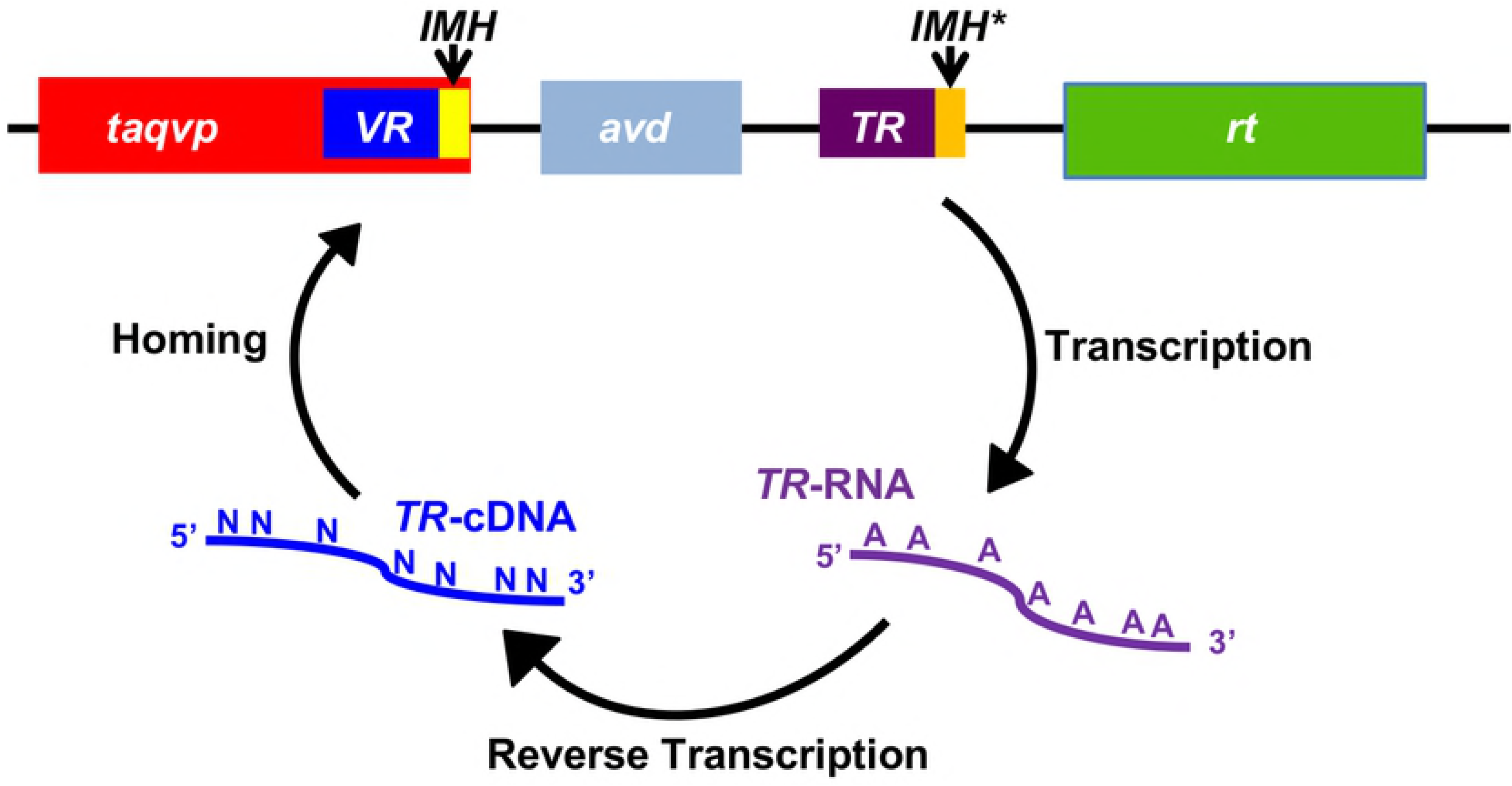
*T. aquaticus* DGR. The T. aquaticus DGR contains a gene encoding a variable protein (*taqvp*). The 3’ end of taqvp contains the variable region (*VR*) and initiation-of-mutagenic homing (IMH) sequence. The DGR also contains an accessory variability determinant (*avd*) gene, followed by an invariant template region (*TR*), which differs from *VR* mainly at adenines. A sequence similar but not identical to the VR IMH occurs at the 3’ end of the *TR* and is called IMH*. Following these elements is a gene encoding a reverse transcriptase (*rt*). TR is transcribed to produce TR-RNA, which is reverse transcribed to produce TR-cDNA, with adenine-specific mutagenesis of the sequence accompanying reverse transcription. Adenine-mutagenized *TR*-cDNA homes to and replaces *VR* to yield a variant TaqVP.

## Results and Discussion

### Structure of TaqVP

TaqVP was overexpressed in *Escherichia coli, purified, and crystallized*. Single-wavelength anomalous dispersion (SAD) data from selenomethionine-labeled crystals of TaqVP were collected and used to determine the structure of TaqVP. The structure was determined and refined to 2.81 Å resolution limit (Table 1), and the entire length of TaqVP, amino acids 1–381, was modeled. TaqVP appeared by gel filtration chromatography to be monomeric in solution and was observed to be monomeric in the crystal (Fig. 2a and 2b).

**Table 1.** Data collection, phasing and refinement statistics.

|  |  |
| --- | --- |
| **Data collection** |  |
| Space group | P 3<sub>2</sub> 2 1 |
| Cell dimensions |  |
| *a*, *b*, *c* (Å) | 155 |
|  | 155 |
|  | 202 |
| $\alpha$, $\beta$, $\gamma$(°) | 90, 90, 120 |
| Wavelength | 0.979 Å |
| Resolution (Å) | 134.27-2.81 (3.29-2.81)<sup>a</sup> |
| *R*<sub>merge</sub> | 0.18 (1.00) |
| *I* / $\sigma_I$ | 12.5 (1.6) |
| Completeness (%) | 100 (100) |
| Redundancy | 7.4 (6.9) |
| cc<sub>1/2</sub> | 0.98 (0.60) |
| **Refinement** |  |
| Resolution (Å) | 80.81-2.81 (3.29-2.81) |
| No. reflections | 132403 (12794) |
| *R*<sub>work</sub> / *R*<sub>free</sub> | 0.18 (0.31)/0.23 (0.33) |
| No. atoms |  |
| Se | 18 |
| Ca | 8 |
| S | 4 |
| O | 2384 |
| N | 2032 |
| C | 7223 |
| H | 90 |
| Average *B*-factors | 60.1 |
| R.m.s deviations |  |
| Bond lengths (Å) | 0.009 |
| Bond angles (°) | 1.22 |
| MolProbity score | 2.1 [98<sup>th</sup>]<sup>b</sup> |
| Ramachandran |  |
| % preferred | 93.7 |
| % allowed | 5.8 |
| % disallowed | 0.5 |
| Clashscore | 7.6 [99<sup>th</sup>]<sup>b</sup> |

<sup>a</sup>Highest resolution bin in parentheses here and other rows.
<sup>b</sup>Percentile in brackets here and other rows.

The structure of TaqVP revealed a single globular CLec-fold domain that strongly resembles the CLec-fold of two previously determined bacterial DGR variable proteins (Fig. 2c and 2d). Specifically, TaqVP closely resembles TvpA (2.5 Å rmsd; 109 Cα; Z = 6.0; 15.8% sequence identity) and Mtd (2.6 Å rmsd; 102 Cα;Z = 5.4, 20.9% sequence identity) [3, 20]. Like TvpA and Mtd, TaqVP has the formylglycine-generating enzyme (FGE) subtype of the CLec-fold and, like these other two proteins, shares structural homology with human FGE (3.2 Å rmsd; 116 Cα; Z = 6.2; 22% sequence identity) [26]. TaqVP is more distantly related to the CLec-fold of the archaeal DGR variable protein AvpA (3.3 Å rmsd; 114 Cα;Z = 3.7, 14% sequence identity), which does not belong to the FGE subtype [22].

**Figure 2.**
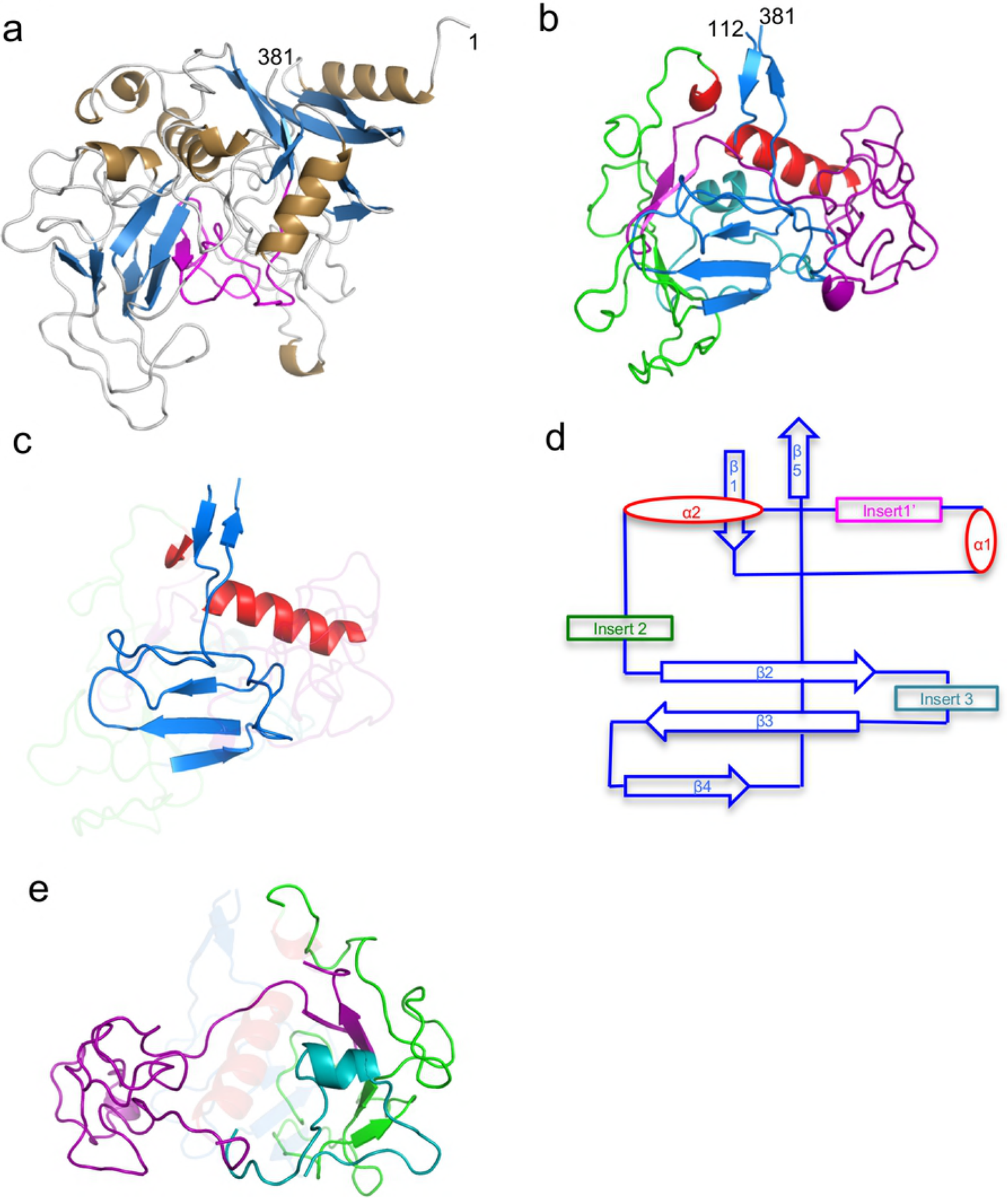
Structure of TaqVP. **a.** TaqVP in ribbon representation (α-helices gold, β-strands blue, loops grey, and VR purple). The amino acid positions of the N- and C-termini of TaqVP are indicated. **b**. TaqVP in ribbon representation with the core elements of the CLec-fold in red (α-helices) and blue (β-strands). Insert 1’, amino acids 129–217, is magenta; insert 2, amino acids 222–296, green; insert 3, amino acids 301–331, teal. **c**. The core elements of the CLec-fold in TaqVP in ribbon representation (α-helices red, β-strands and loops blue). The inserts are ghosted. **d**. Topology diagram of the CLec-fold in TaqVP. **e**. Inserts of TaqVP in ribbon representation, with core elements of the CLec-fold are ghosted (color coding same as in panel b).

The FGE-type CLec domain of TaqVP begins at amino acid 112 and extends to its C-terminal amino acid 381. Preceding amino acid 112 is a region composed of short β-strands and α-helices that wraps around the N-terminus of the CLec domain. The characteristic CLec-fold features, including the N- and Ctermini forming anti-parallel β-strands (β1 and β5) and two roughly perpendicular α–helices (α1 and α2), are found in TaqVP. The second classic feature of the CLec fold in DGR variable proteins is a four-stranded, anti-parallel β-sheet (β2 β3 β4 β4’). In TaqVP the β4’ strand is replaced by a loop. The sequence between the β3 and β5 strands forms the putative ligand-binding site. TaqVP also has short segments inserted between the secondary elements of the CLec fold (Fig. 2e), as also seen in Mtd, TvpA, and AvpA.

### Variable Region

The variable region of TaqVP (amino acids 341–381) is located at the very C-terminus of the protein, as in Mtd and TvpA. The TaqVP VR is 40 amino acids in length — between the 45-amino acid VRs of Mtd and TvpA and the 23-amino acid VR of AvpA. As shown previously for all structurally characterized DGR variable proteins, the nine variable amino acids in TaqVP are solvent exposed and form a potential binding site (Fig. 3a and 3b). Variable hydrophobic and hydrophilic amino acids are segregated from each other at this site (Fig. 3b). TaqVP has two nonvariable aromatic amino acids (W349 and W371) within the binding site, which might provide a constant hydrophobic contact to ligands. Similarly, Mtd, TvpA, and AvpA have one or two nonvariable aromatic acids within their ligand-binding sites.

**Figure 3.**
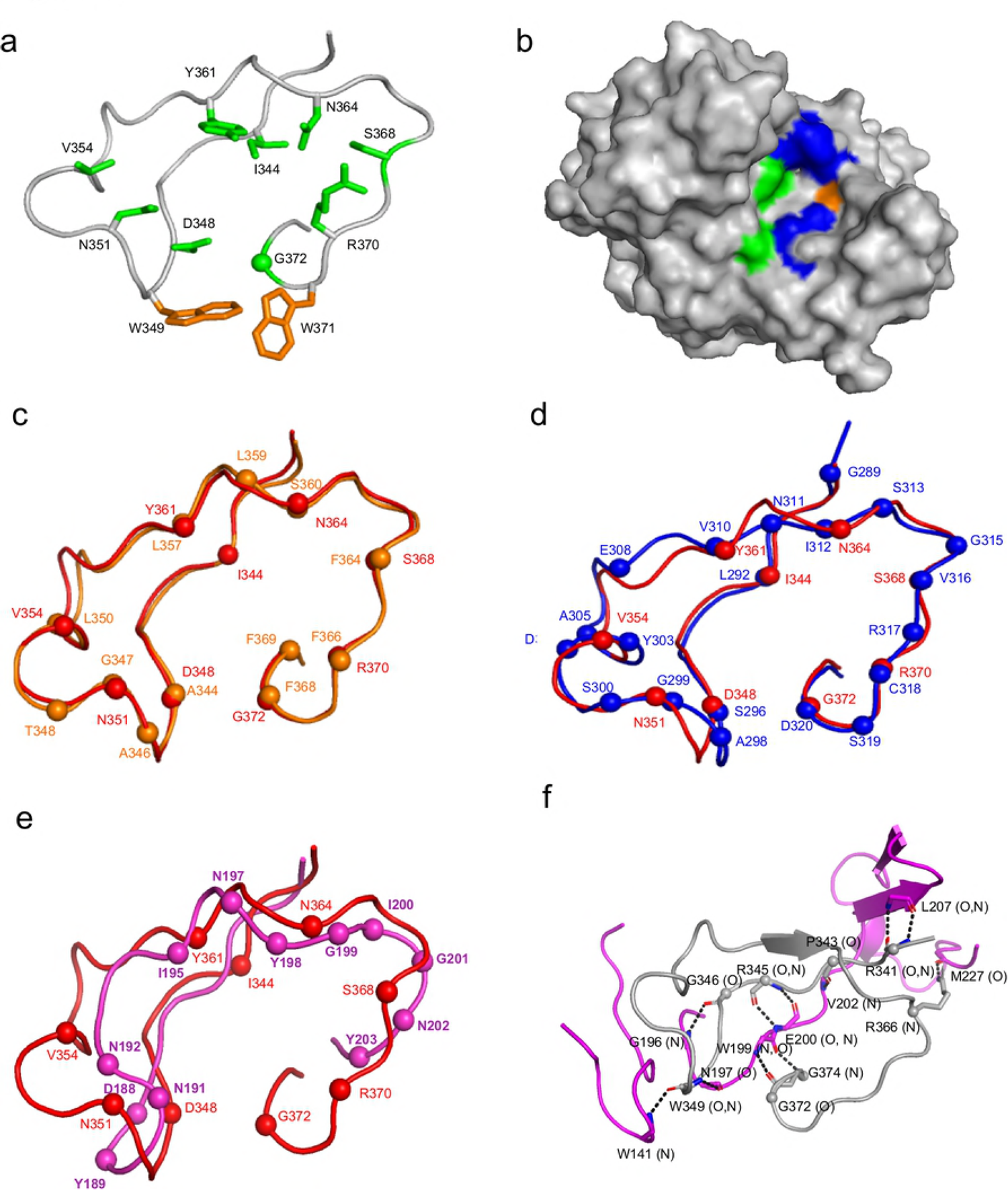
Variable region of TaqVP. **a.** VR of TaqVP in ribbon representation. The main chain is in gray, side chains of variable amino acids are in green (sphere is glycine), and nonvariable aromatic amino acids are in orange. **b**. Surface representation of TaqVP, with the VR facing the viewer. Variable hydrophobic amino acids (I, V, and Y) are green, variable hydrophilic amino acids (S, N, D, and R) blue, and variable glycine pale orange. **c**. Superposition of the VR of TaqVP (red) and Mtd-P1 (orange) in Cα representation. The spheres represent the position of variable amino acids in each protein. **d**. Superposition of the VR of TaqVP (red) and TvpA (blue) in Cαcrepresentation. **e**. Superposition of the VR of TaqVP (red) and AvpA (magenta) in Cα representation. **f**. Stabilization of the main chain of VR (gray, Cα of variable amino acids indicated by spheres) by insert 1’ (magenta) in Cαcrepresentation(c7ashedclinec indicates hydrogen bonds.

Remarkably, the TaqVP variable region and those of Mtd and TvpA are nearly identical in structure, despite their weak sequence relationship (rmsd 0.46 Å, 26 Cα, p<0.001, 28.9% sequence identity; and rmsd 1.10 Å, 30 Cα, p<0.001, 23.1% sequence identity, respectively) (Figs. 3c and 3d). However, the TaqVP variable region does not share significant structural similarity with the variable region of the more distantly related AvpA (rmsd, 2.28 Å; 22 Cα; p>0.1) (Fig. 3e). The nine variable amino acids in TaqVP VR support a potential diversity through adenine-mutagenesis of 3 × 10^9^. Seven of the 9 variable amino acids are encoded by AAY codons in *TR* (D348, V354, Y361, N364, S368, and R370 by AAC and N351 by AAT), enabling substitution by 14 other amino acids at each of these positions. The remaining two variable amino acids are encoded by an ATC (I344) or AGT (to G372) codon, which enables substitution by three other amino acids. All nine of the TaqVP variable amino acids have structural equivalents in TvpA, while eight of the nine have structural equivalents in Mtd (Figs. 3c and 3d). The structural similarity among these VRs suggests that a composite VR having maximal diversity may be designed from these.

The last seven amino acids (375–381) of TaqVP VR are invariant and located along the β5-strand, as in Mtd and TvpA. The corresponding DNA sequence is likely to function as the initiation of mutagenic homing (IMH) element, a critical component of DGRs that along with the IMH* element in *TR* defines the directionality of cDNA transfer [18] (Fig. 1).

### Inserts

TaqVP has three inserts within the core of the CLec-fold. Insert 1’ (amino acids 129–217) is located between α1 and α2; an equivalent to insert 1’ occurs in TvpA. Insert 1’ in TaqVP appears to stabilize the VR through hydrogen bonding (Fig. 3f), whereas insert 1’ in TvpA interacts with helix α1 instead. Insert 2 of TaqVP is between α2 and β2 (amino acids 222–296). An equivalent occurs in all DGR variable proteins structurally characterized to date, but unlike these others, insert 2 in TaqVP does not interact with the VR. Insert 3 (amino acids 301–331) is between β2 and β3 of the CLec-domain and is composed of loops and a short α-helix; an equivalent for insert 3 occurs in AvpA.

### Thermal Stability

The thermal stability of TaqVP was determined by monitoring its secondary structure as a function of temperature by circular dichroism. We observed that TaqVP started to unfold at ∼70 °C and was completely denatured at ∼90 °C, as monitored by the loss of ellipticity (Fig. 4a). The thermal unfolding of TaqVP was irreversible. AvpA has previously been studied for its thermal stability and it was found that this protein also adopts a thermostable fold, starting to unfold at ∼65 °C and becoming completely denatured at ∼80 °C [10]. We also determined the thermal stabilities of Mtd-P1 and TvpA. *Bordetella* bacteriophage Mtd-P1, which unlike the other DGR variable proteins studied here has multiple domains, was found to start irreversibly unfolding at ∼70 °C and was completely denatured at ∼80 °C (Fig. 4b). TvpA began unfolding at ∼50 °C and was completely denatured at ∼70 °C (Fig. 4c). For TvpA, the ellipticity became more negative upon unfolding. This shift to more negative ellipticity upon unfolding has been seen previously in the unfolding of dihydrofolate reductase due to the involvement of W47 and W74 forming an exciton pair [27]. TvpA has a structurally similar pairing of tryptophans (W138 and W263) that could form an exciton pair.

**Figure 4.**
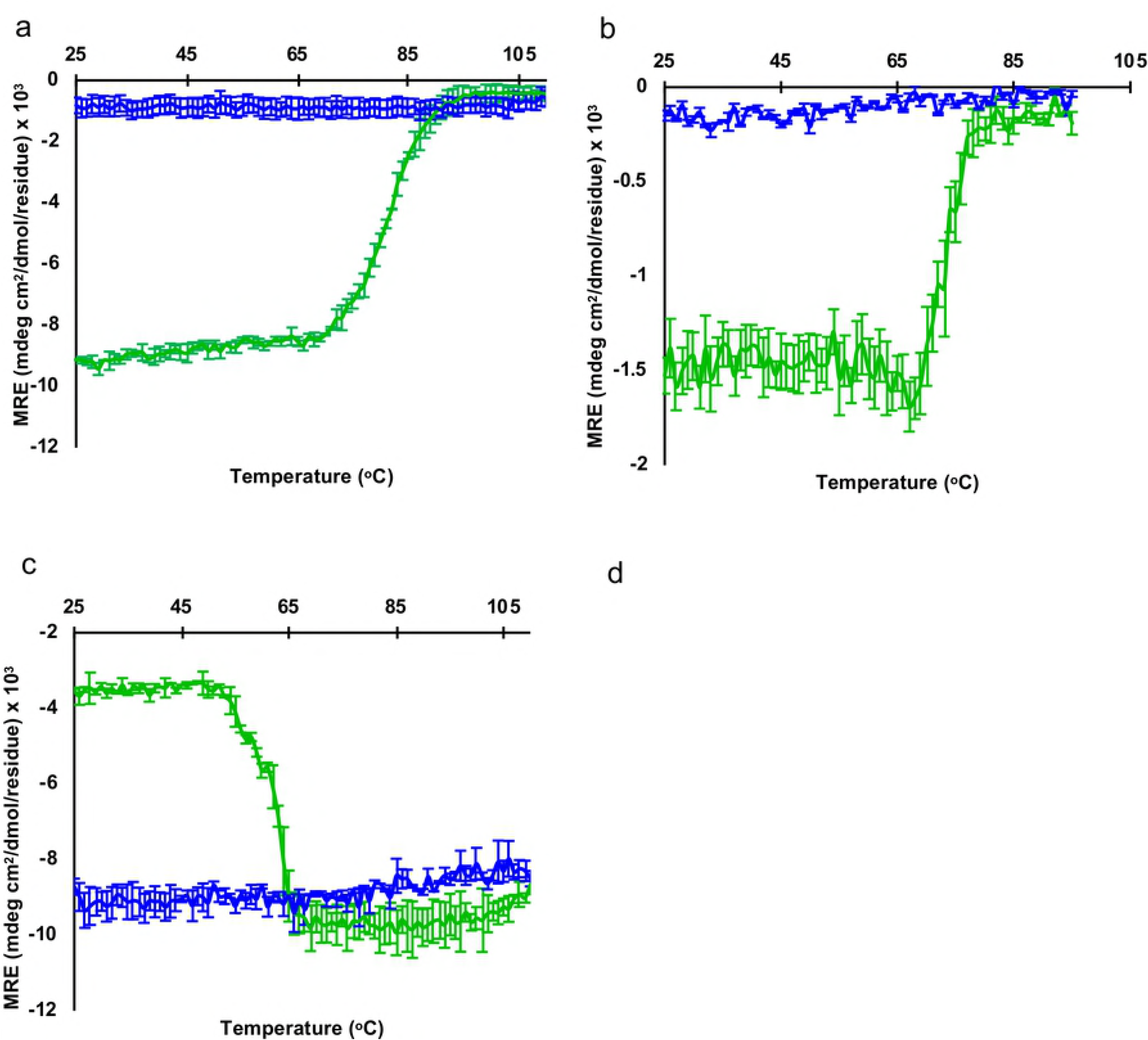
Thermal stability of DGR variable proteins. Circular dichroism signal (mean residue ellipticity, MRE) at 216 nm for the transition from 25 to 110 °C (green), and transition from 110 to 25 °C (blue) for TaqVP (a), Mtd-P1 (b), and TvpA (c). Standard deviations from three separate experiments are shown.

These results suggest that thermostability is shared by several of the structurally characterized CLec-fold DGR variable proteins (i.e., Mtd, AvpA, and TaqVP), and that any of these may provide an advantageous scaffold for molecular display applications.

## Methods

### TaqVP expression and purification

The coding sequence of TaqVP (accession no. CP010822.1) was synthesized with codons optimized for expression in *Escherichia coli* (GENEWIZ, Inc.) and cloned into a modified pET28b expression vector encoding an N-terminal His-tag followed by a PreScission protease cleavage site. The integrity of the construct was confirmed by DNA sequencing. TaqVP was expressed in *E. coli* BL21-Gold (DE3). Bacteria were grown with shaking at 37 °C to an OD_600_ of 0.6–0.8 and then cooled to room temperature, followed by induction with 0.5 mM isopropyl β-D-1-thiogalactopyranoside. Bacteria were grown with shaking at room temperature for 4 h further, then harvested by centrifugation (30 min, 5,000 × g, 4 °C); the bacterial pellet was frozen at −80 °C.

Cells were thawed and resuspended in buffer A (250 mM NaCl, 50 mM Tris, pH 8, and 5 mM β-mercaptoethanol; 20 ml/L of bacterial culture) supplemented with 1 mM phenylmethylsulfonyl fluoride. Bacteria were lysed using an Emulsiflex (Manufacturer), and the lysate was centrifuged (30 min, 35,000 × g, 4 °C). The supernatant was incubated at 55 °C for 10 min, and the sample was centrifuged (30 min, 35,000 × g, 4 °C). The supernatant was then applied to a column containing His-Select Nickel affinity gel (Sigma, 1 ml of resin per 20 ml of bacterial lysate), which had been equilibrated with buffer A. The column was washed with 10 column volumes of buffer B (250 mM NaCl, 20 mM Tris, pH 8, and 5 mM β-mercaptoethanol) containing 20 mM imidazole, and the TaqVP was eluted with buffer B containing 250 mM imidazole. The His-tag was removed by PreScission protease cleavage (1:50 mass ratio TaqVP:protease) overnight at 4 °C. Cleaved TaqVP was separated from the non-cleaved protein by applying the sample to a His-Select Nickel affinity gel column (Sigma) and collecting the flow-through. TaqVP was further purified by gel filtration chromatography (Superdex 75) in 150 mM NaCl, 20 mM Tris, pH 8, and 1 mM dithiothreitol.

### Crystallization and structure determination

Selenomethionine (SeMet)-substituted TaqVP was expressed by culturing *E. coli* in synthetic minimal media supplemented with 200 mg/L L(+)− selenomethionine (Sigma) [28]. Purified SeMet-labeled TaqVP was concentrated to 50 mg/mL by ultrafiltration (10 kDa MWCO Amicon, Millipore); the concentration of TaqVP was determined using a calculated molar extinction coefficient at 280 nm of 80,900M^−1^cm^−1^.

Crystals of SeMet-labeled TaqVP were grown by the hanging drop method at 20 °C by mixing 1 μL of TaqVP (50 mg/mL) and 1 μL of 15 % (w/v) 2-methyl-2,4-pentanediol, 20 mM CaCl_2_, 100 mM sodium acetate, pH 4.6. Crystals were cryoprotected by soaking in the precipitant solution supplemented with 20% glycerol. Single-wavelength anomalous dispersion (SAD) data were collected at Advanced Photon Source (Argonne, IL) beamline 24-ID-E. Diffraction data were indexed, integrated, and scaled with MOSFLM [29–31]. Se sites were located from SAD data of SeMet-labeled TaqVP, and initial phases were determined using SOLVE [32]. All five methionines (M1, M24, M112, M227, and M320) were located, and the asymmetric unit was found to contain four molecules of TaqVP.

An initial model of TaqVP was built by automatic means using Autobuild (within Phenix). A total of 75 iterative rounds of manual model building and maximum likelihood refinement were carried out with Refine (within Phenix) using default parameters, with each refinement step consisting of 3 cycles [33, 34]. Each refinement step included manual model rebuilding with COOT, guided by σ_A_-weighted 2mF_o_-DF_c_ and mF_o_-DF_c_ difference maps [35]. One round of TLS parameterization with default settings was then used, followed by the addition of water, calcium, and acetate ions into ≥3σ mF_o_-DF_c_ density. Structure validation was carried out with Molprobity [36], and molecular figures were generated with PyMOL (http://www.pymol.org/). The crystal structure and structure factors have been deposited to the Protein Data Bank (accession no. 5VF4).

### Structural alignment of VR and equivalent regions

The structure of the VR of TaqVP (amino acids 341–374) was compared to that of the VR of Mtd (amino acids 337–381), TvpA (amino acids 285–329), and AvpA (amino acids 181–210) using FATCAT [37].

### CD Spectroscopy

CD spectra of Taq VP, Mtd-P1, and TvpA were collected between 195 and 260 nm at 24 °C with 1-nm intervals, and temperature denaturation scan was carried out between 25 °C and 110 °C at 216 nm with 1° intervals, using a quartz cell with a 1-mm path length on a model 202 spectrometer (Aviv Instruments) equipped with thermoelectric temperature control. Measurements were collected for Taq VP, Mtd-P1, and TvpA samples at 0.4 mg/mL, 0.3 mg/mL and 0.3 mg/mL, respectively. Protein samples were in 40 mM NaF, 10 mM NaP*i* buffer, pH 7.5. The CD signal from buffer alone was subtracted from the data before conversion to mean residue ellipticity.

